# Calibrated feedback illumination for precise conventional fluorescence and PALM imaging applications

**DOI:** 10.1101/718981

**Authors:** A. Mancebo, L. DeMars, C. T. Ertsgaard, E. M. Puchner

## Abstract

Spatial light modulation using cost efficient digital mirror arrays (DMA) is finding broad applications in fluorescence microscopy due to the reduction of phototoxicity and bleaching and the ability to manipulate proteins in optogenetic experiments. However, the precise calibration of DMAs and their application to single-molecule localization microscopy (SMLM) remained a challenge because of non-linear distortions between the DMA and camera coordinate system caused by optical components. Here we develop a fast and easy to implement calibration procedure that determines these distortions by means of an optical feedback and matches the DMA and camera coordinate system with ~50 nm precision. As a result, a region from a fluorescence image can be selected with a higher precision for illumination compared to manual alignment of the DMA. We first demonstrate the application of our precisely calibrated light modulation by performing a proof-of concept fluorescence recovery after photobleaching experiment with the endoplasmic reticulum-localized protein IRE1 fused to GFP. Next, we develop a spatial feedback photoactivation approach for SMLM in which only regions of the cell are selected for photoactivation that contain photoactivatable fluorescent proteins. The reduced exposure of the cells to 405 nm light increases the possible imaging time by 44% until phototoxic effects cause a dominant fluorescence background and a change in the cell’s morphology. As a result, the mean number of reliable single molecule localizations is also significantly increased by 28%. Since the localization precision and the ability for single molecule tracking is not altered compared to traditional photoactivation of the entire field of view, spatial feedback photoactivation significantly improves the quality of SMLM images and the precision of single molecule tracking. Our calibration method therefore lays the foundation for improved SMLM with active feedback photoactivation far beyond the applications in this work.

**Statement of significance:** Actively patterned illumination in fluorescence microscopy can reduce bleaching and phototoxicity as well as actively manipulate proteins in optogenetic applications. Matching the coordinate system of the camera and the light patterning device such as digital mirror arrays (DMA) remains a challenge. We developed a fast and easy calibration procedure that determines and corrects for the transformation between the camera and DMA coordinate system with ~50 nm precision. Using this approach, we develop spatial feedback photoactivation for Single Molecule Localization Microscopy (SMLM) to photoswitch only intracellular regions containing photoswitchable fluorophores. Our results show a 44% improvement in the possible data acquisition time before phototoxic effects become detectable and a 28% increase in detected localizations. Spatial feedback photoactivation thus significantly improves SMLM experiments.

## Introduction

Fluorescence microscopy techniques are continually advancing our understanding of cellular processes through the specific labeling and visualization of biomolecules within living cells. The quantitative analysis of this data provides valuable information about the complex spatial organization, dynamics and interactions of biomolecules, which are essential for all fundamental biological processes. In traditional confocal and epi- fluorescence microscopy applications, the excitation power is kept constant across the field of view. The resulting fluorescence intensity therefore correlates with the density of labeled molecules within a diffraction limited volume. Over the decades, however, many advanced imaging modes have been developed that rely on either excitation light being focused to a region of interest (ROI) within the sample or spatially-patterned illumination across the sample. Fluorescence fluctuation spectroscopy, for instance, makes use of a locally excited region to measure the intensity fluctuation of fluorophores that diffuse in and out of a focused laser beam. From the recorded data the concentration of labeled proteins, their diffusion coefficients (1) and their oligomeric states (2) can be quantified. Fluorescence recovery after photobleaching (FRAP) applications utilize a small selected volume within the sample that is excited to locally deplete its fluorophores through bleaching. The diffusion coefficient and the immobile fraction of labeled biomolecules can then be determined based from the time and the fraction of fluorescence recovery within the ROI (3).

Patterning the excitation light within the sample plane has also found applications in active illumination fluorescence microscopy for long-term measurements. By actively regulating the excitation to keep a fixed detection power across the field of view, detector saturation is avoided while maintaining sensitivity for weak signals (4, 5). The active patterning of light in the sample plane is not only used for imaging but also for manipulating proteins in optogenetic applications. By locally activating light sensitive ion channels or the interaction of regulatory proteins within a cell, important information about spatial aspects of cell signaling has been obtained (6–8).

While all fluorescence microscopy techniques employing spatially patterned light excitation made significant and valuable contributions to our understanding of cellular processes, they are all hampered by the diffraction-limited readout of the fluorescence signals. This limited resolution prevents resolving the myriad of intracellular structures and protein assemblies that are much smaller than the optical diffraction limit of ~250 nm. The advent of fluorescence super-resolution microscopy techniques overcame these limitations and allows us to resolve intracellular structures with ~20 nm resolution (9–12) and to quantify the biomolecules they contain (13–15). Single-molecule localization microscopy (SMLM) methods, termed stochastic optical reconstruction microscopy (STORM) (9) and photoactivated localization microscopy (PALM) (10), are based on activating only a sparse subset of photoactivatable fluorophores in each frame either by photoswitching or by exploiting intrinsic dark states of the fluorophores (9, 10, 16). Similarly, fluorogen activating proteins have been used to create spatially separated fluorescent bursts of transiently binding fluorogens (17, 18). The center location of individual fluorophores can be then be determined with high precision by fitting each point-spread function in each frame. A super-resolution image is then built by combining the single-molecule localizations from numerous data acquisition frames.

In this work we present a simple computer-based calibration approach for a versatile, high-precision and low-cost setup that combines for the first time the advantages of actively patterned feedback illumination with SMLM imaging. We first describe the construction of an excitation path comprising a Digital Mirror Array (DMA) that can be added to existing fluorescence microscope setups. While DMAs have been used in the past in custom-made and commercial setups for patterned illumination (19–22) a persistent problem for high-precision applications is the nonlinear distortion between the DMA and the camera coordinate system caused by optical components. Our custom computer-based feedback calibration determines and corrects for these distortions. As a result, the DMA pixels are matched to the camera pixels with a median deviation of 52 nm, which is within the resolution range of fluorescence super-resolution microscopy techniques and cannot be obtained by manual alignment. We demonstrate the versatility of this approach for patterned illumination applications by performing a proof of concept FRAP experiment. We further combine, for the first time, active feedback illumination with SMLM by photoactivating only those regions within the cell where photoactivatable fluorescent proteins (PAFPs) are located. We demonstrate that patterned photoactivation significantly reduces background fluorescence caused by phototoxicity, increases the possible imaging time nearly 44% and provides a 28% improvement in the number of detected single-molecule localizations compared to traditional photoactivation of the entire field of view. This increase in localizations improves the quality of super-resolution images as well as the accuracy of single molecule tracking data. Our calibration method therefore presents a versatile, precise and cheap option for microscopy applications with actively patterned illumination and lays the foundation for improved SMLM with active feedback illumination far beyond the demonstrated applications in this work.

## Materials and methods

### Experimental Setup

All conventional and super-resolution images were recorded on an inverted microscope (Ti-E; Nikon, Minato, Tokyo, Japan) with a Perfect Focus System to compensate for sample drift in the z-direction. All movies were recorded on an electron-multiplying charge-coupled device (EMCCD) camera (iXon 897 Ultra DU-897U; Andor, Belfast, United Kingdom). The camera was cooled down to −70°C. The EMCCD gain was set to 30 for all experiments except the bleaching portion of FRAP experiments for which it was set to 0 (no EMCCD gain).

Lasers (405 nm, 488 nm, 561 nm, OBIS-CW; Coherent, Santa Clara, CA) were combined with dichroic mirrors, aligned, expanded, and focused to the back focal plane of the objective (Nikon-CFI Apo 100x Oil immersion NA 1.49). The lasers’ intensities were controlled by a computer via USB and were digitally-modulated using a NI-DAQ board (PCI-6221; National Instruments, Austin, TX) connected to the computer with programmed shutter sequences. The NI-DAQ board synchronized the lasers to the camera frame duration.

All illumination patterning was done with a DMA (DLP® LightCrafter 4500™; Texas Instruments, Dallas, TX) consisting of 912×1,140 square pixels arranged in a diamond configuration, with a refresh rate of 60 Hz. Each pixel has a side length of 7.639 μm, which corresponds to 114 nm in the sample plane. To specify the illumination, a binary mask is sent to the DMA using custom software written in Python. While in principle a binary mask can be created by thresholding a fluorescence image, we hand-drew the regions based on the fluorescence image for more control. The transformation is then applied to the pattern before sending it to the DMA to account for misalignments and distortions caused by the optical components between the DMA and camera. For PALM experiments, the expanded 405 nm laser was directed onto the DMA by mirrors with an angle of incidence of 24° to the DMA surface normal. This angle is the blaze angle of the DMA and is required to maximize diffraction efficiency and for the reflection to be parallel to the DMA surface normal. The diffracted light from the DMA was captured by a 4F system consisting of a lens of 1 in diameter and focal length of 50.2 mm, a mirror mounted at an angle of 45° and a lens of 2 in diameter and focal length 150 mm. This 4F system serves two purposes: to capture the diverging orders of the DMA near the source, maximizing power transmission, and to form the DMA image in the focal plane of the microscope lens. A longpass dichroic filter (ZT405rdc; Chroma, Bellows Falls, VT) was used to combine the patterned light of the 405 nm laser with the 488 nm and 561 nm beams before they are focused by a lens of 2 in diameter and focal length of 400 mm to the back focal plane of the microscope objective (see Fig. S1). For FRAP experiments, a longpass dichroic filter (ZT488rdc; Chroma, Bellows Falls, VT) was used instead to direct the patterned light of the 488 nm laser into the microscope.

For two-color imaging of red (595 nm) and green (525 nm) fluorescence, the fluorescence emission was split by a dichroic longpass beamsplitter (T562lpxr BS; Chroma, Bellows Falls, VT). The emission in each channel was further filtered by bandpass filters: ET525/50m (Chroma, Bellows Falls, VT) in the green channel and ET595/50m (Chroma, Bellows Falls, VT) in the red channel.

To estimate the power density of the lasers at the sample plane, we measured the intensity after all neutral density filters but before the beam expander with a power sensor (S130C; Thorlabs, Newton, NJ) and divided by the illumination area in the sample plane. The typical power density for 561 nm excitation of single molecules was 428 W/cm^2^ and for conventional 488 nm excitation 0.15-0.45 W/cm^2^. To estimate the power density of patterned light, we measured the emission of a fluorescein dye sample with 405 nm excitation. The DMA power density was then calculated by multiplying the power density without the DMA by the mean ratio of the emission intensities with and without the DMA. The typical power density of patterned 405 nm activation (which was gradually increased during each PALM experiment to maintain the activation rate) was 3.6 mW/cm^2^-177 W/cm^2^. The power density of patterned 488 nm excitation for bleaching in FRAP was 89 W/cm^2^. The power density for initial fluorescence and fluorescence recovery measurements in FRAP was 1.7 W/cm^2^.

### Sample preparation

#### Concanavalin A (ConA)

Cells were immobilized to the chambered coverglass surface by using a solution of ConA in deionized water (0.8 mg/mL; Sigma-Aldrich, St. Louis, MO) which was incubated on the coverglass surface. After at least 30 minutes, excess ConA was withdrawn and the slide was rinsed three times with deionized water prior to plating the cells. Diluted cells were allowed to settle and bind to the ConA on the surface for at least 20 minutes before imaging.

#### DMA Calibration

Calibration was performed by projecting the calibration pattern with 405 nm excitation onto a coverslip with a solution of fluorescein dye (10-20 μL of 130 μM; Sigma-Aldrich, St. Louis, MO) .

#### FRAP

FRAP experiments were performed using a W303 MATa (MATa ura3 trp1 leu2 his3 ade2 can1, Horizon-Dharmacon, YSC1058) *S. cerevisiae* strain. The ade2 mutation in the purchased strain was repaired by PCR-mediated transformation. Wild-type ADE2 was amplified from genomic DNA, RB201 (W303 MATa, trp1, leu2, ura3, his3, can1R, ADE2) with Phusion PCR (NEB) using the forward primer (ATGGATTCTAGAACAGTTGGTATATTGGGAGGGGGACAA) and the reverse primer (TTACTTGTTTTCTAGATAAGCTTCGTAACCGACAGTTTCTAACTT). To label IRE1 with GFP (IRE1-GFP-IRE1), the native IRE1 gene was knocked out by a PCR-mediated gene deletion using the pFA6a-kanMX6 template (23) and forward primer: CCTTCATACACATTAAAAAAACAGCATATCTGAGGAATTAATATTTTAGCACTTTGAAAAtacgctgcaggtcgacgg and reverse primer: ATGATCAAAGTAACATTAATGCAATAATCAACCAAGAAGAAGCAGAGGGGCATGAACATGcatcgatgaattcgagctcg. The knockout was confirmed by colony PCR. Next, the IRE1 promoter (1346 base pairs upstream the start codon) and the IRE1 gene until the Juxtamembrane position, which is a tagging site that does not interfere with the function of IRE1 (24), was PCR amplified from W303 genome using forward primer: TCGTA GGGCCC GTCGTGCTATGTTGAGAAACGA to introduce the ApaI restriction site and the reverse primer: ACATTCTCGAGACCAGATCCAATTTTGGATAATAATACATA at the Juxtamembrane position to introduce the XhoI restriction site. yeGFP was PCR amplified using the forward primer: TCGTA CTCGAGatgtctaaaggtgaagaattattca to introduce the XhoI restriction site and the reverse primer: AATGTGGATCCtttgtacaattcatccatacc to introduce the BamHI restriction site. The remaining IRE1 gene up to the stop codon was PCR amplified with the forward primer TCGTAGGATCCGGATTTATGCCTGAAAAGGAAAT to introduce and additional BamHI restriction site and reverse primer TCGTAGCGGCCGCTTATGAATACAAAAATTCACGTAAA to introduce a NotI restriction site after the stop codon. PCR products were assembled in pNH605 yeast integration plasmid that targets the LEU2 locus (25) and transformed in the W303 IRE1 knockout strain. Integration was confirmed by colony PCR and fluorescence imaging.

The cells were inoculated in synthetic complete dextrose (SCD) medium to an optical density of 0.28 and grown overnight in a shaking incubator at 270 rpm and 30 °C. The cells were diluted to an optical density (OD) of 0.29 in the morning and grown in the shaking incubator under the same conditions for 3.5 hours until and OD of 0.78 was reached. The cells were again diluted to an OD of 0.31 and after 2.5 hours, the cells reached an OD of 0.72. The cells were then diluted 1:7 to achieve an OD of 0.1 and 210 μL were plated and allowed to settle for at least 20 minutes.

#### PALM

PALM experiments were performed using a W303 *S. cerevisiae* strain (pSte5_mEos2-3x_PH2x) (26) with a triple-repeat of mEos2 fused to the plextrin homology (PH) domain of Plc(delta1), which localizes to the plasma membrane. The construction of this yeast strain was previously published in reference (26) and for this study we replaced the weak pINO promoter with 511 bases of the stronger pSte5 promoter (W303 genome). The cells were inoculated in SCD medium to an optical density of 0.13 and grown overnight in a shaking incubator at 270 rpm and 30 °C. The cells were diluted to an OD of 0.23 in the morning and grown in the shaking incubator under the same conditions for 4 hours to OD 0.72. To locate the plasma membrane for patterned photoactivation during PALM imaging, the yeast cell wall was stained with ConA-CF488M (Biotium, Fremont, CA) which is a 488 nm-excitable dye conjugated to ConA. The cells were incubated at an OD 0.1 for 30 minutes in SCD containing 33 μg/mL of ConA-CF488M in a shaking incubator at 270 rpm and 30 °C.

### Data acquisition and analysis

#### DMA Calibration

To precisely map the DMA pixels to the camera pixels, a 4×4 grid of spots was sent to the DMA and the reflected 405 nm laser was projected onto a sample of fluorescein dye. The resulting emission spots with 1.83 μm diameter and 9.53 μm center-to-center spacing were recorded at 5 Hz for 50 frames and the frames were averaged to reduce camera noise. Both the microscope image and the projected image were processed by the software, which detects the spots’ coordinates (Fig. 1 A and S2). This custom software is written in Python and leverages the OpenCV (27) and scikit-image (28) libraries for computer vision and image processing. The software determines the centroid of each spot by blob detection in the microscope image. The detected spots are subsequently fit to Gaussians within a window of twice the estimated spot size. The software for warping and sending images to a DMA and for determining the transformation between a projected image and its corresponding microscope image will be freely available for download on GitHub.

**Figure 1:**
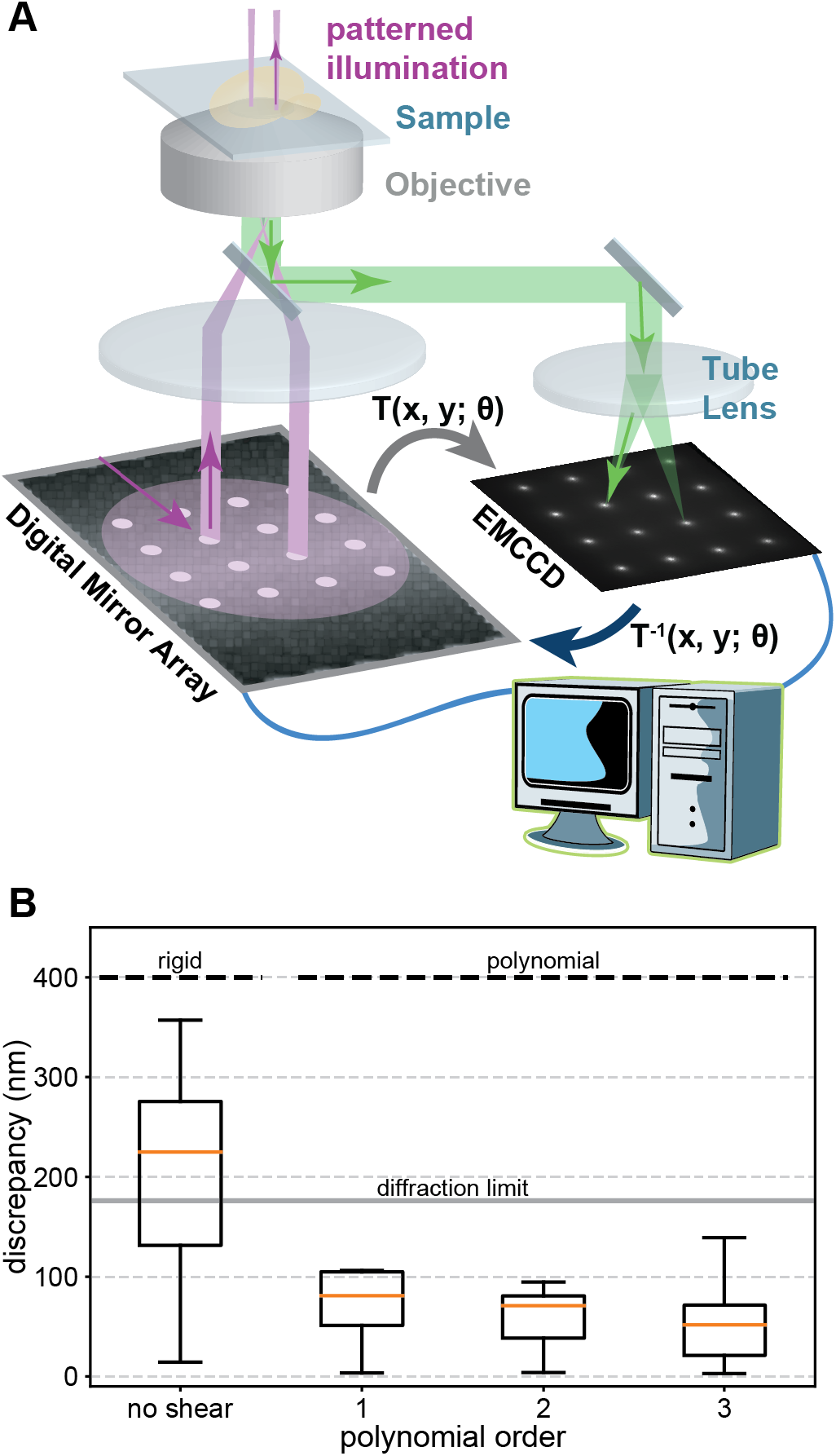
Schematics of spatial feedback illumination setup and calibration. (A) A digital mirror array is mounted in the excitation path of a PALM microscope. Pixels in the on state reflect laser light, which gets projected onto the back focal plane of the microscope objective (purple, 4F lens system not shown for clarity). The fluorescence (green) of the calibration patter of 4×4 spots is recorded with the camera and the transformation T between the DMA and camera coordinate system is determined with a computer to correct for rigid and higher order polynomial distortions caused by optical components (39, 40). (B) Varying degrees of freedom are included in the inverse transformation T^−1^ of the calibration pattern to correct for distortions. (Left) Rigid degrees of freedom simulating a perfect manual alignment result in a significant median discrepancy above the theoretical diffraction limit. (Right) Including higher order polynomials in T^−1^ results in an accurate matching of the DMA and camera coordinate system with a median discrepancy of 52 nm. Boxes extend from the 25^th^ to 75^th^ percentiles (interquartile range, IQR). Medians are shown as orange horizontal lines. Whiskers extend to the farthest data point within 1.5 IQR below and above the 25^th^ and 75^th^ percentiles, respectively.

The transformation between the centroids of the spots in the projected image and the microscope image are determined by sorting the coordinates in two dimensions and finding the nearest neighboring pixel. This sorting scheme is robust under small rotations that don’t cause rows of the pattern to overlap vertically with other rows, i.e., they can be separated vertically. After sorting, one set of coordinates is fit to the other by a 3^rd^-order polynomial of the form shown in Eq. 1

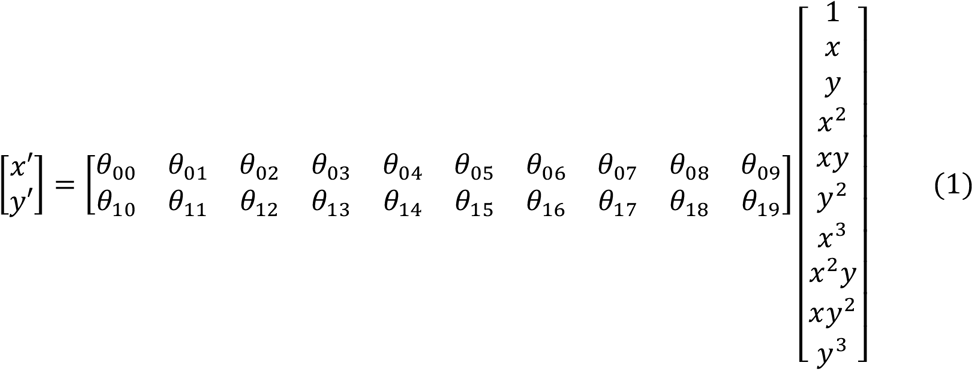

where (*x*, *y*) and (*x*′, *y*′) are the source and destination coordinates, respectively, and {θ_*ij*_ | 0 ≤ *i* ≤ 9 and *j* = 0,1} are the weights for each term in the polynomial transformation that are determined during fitting. Since we compensate for distortions by using the inverse transformation, the source and destination correspond to the camera and DMA, respectively. To verify the accuracy of the transformation and to estimate its precision, we warped the original calibration pattern by applying the inverse transformation to it and sent the warped calibration pattern to the DMA. The warped calibration pattern was projected onto the dye sample, the emission was recorded, and the original calibration pattern and microscope image were processed as previously described in the DMA calibration section (see Fig. S2). The median distance between corresponding points of the original calibration pattern and the camera image was 52 nm, which is a precision well below the theoretical diffraction limit (~200 nm) (Fig. 1 B).

#### FRAP

A 488 nm laser at a power density of 1.7 W/cm^2^ was used to excite GFP in the full field of view for 2 seconds (Fig. 2 A, left). The averaged frames over this time were used to select regions of the ER in each cell to generate a DMA mask for bleaching (Fig. 2 A, second from left). The selected spots were bleached for 60s at a power density of 89 W/cm^2^ (Fig. 2 A, center). To minimize bleaching while measuring the fluorescence recovery, the full field of view was subsequently illuminated with a 488 nm laser pulse with a duration of 0.2 seconds repeated every second at a power density of 1.7 W/cm^2^. To determine the region of interest for measuring fluorescence recovery, a subset of the frames during bleaching were averaged in time and a maximum entropy threshold (29) was applied to the fluorescence image in Fiji. The intersection of the resulting area with the DMA mask was used to measure bleaching and recovery of fluorescence (Fig. 2 A, second from right and right).

**Figure 2:**
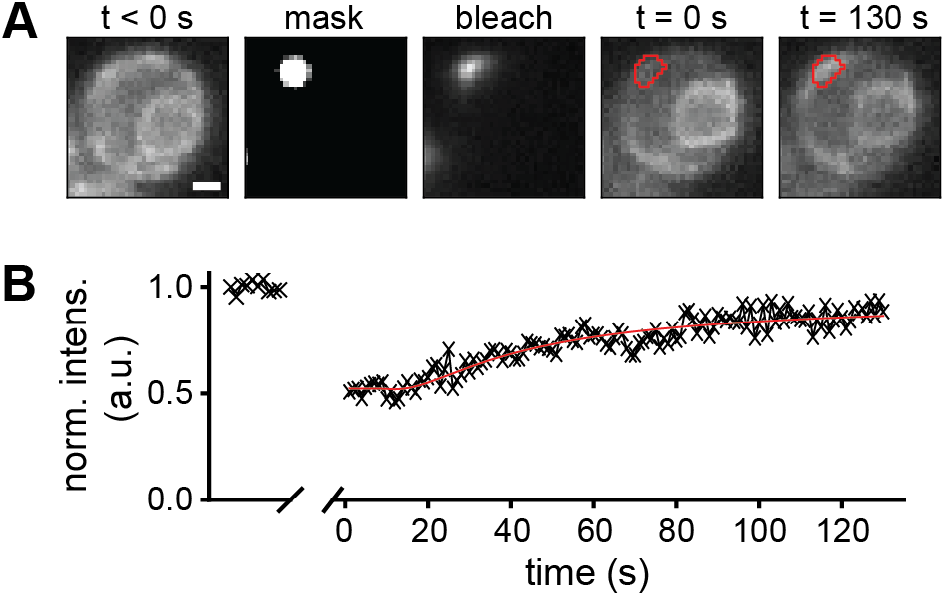
Proof of concept FRAP experiment. (A, left) Fluorescence image of IRE1-GFP before bleaching. (A, second from left) Selected spot for bleaching IRE1-GFP with 488 nm light. (A, center) Fluorescence of IRE1-GFP during bleaching. (A, second from right) Fluorescence image of IRE1-GFP after bleaching of the selected spot (red outline). (A, right) Fluorescence image of IRE1-GFP after fluorescence recovery of the selected spot (red outline) (B) Normalized bleach-corrected intensity of the IRE1-GFP emission within the bleached region outlined in red. Scale bar: 1 μm.

To correct for overall bleaching of the sample during the recovery measurement, which was measured to be less than 5%, each frame was normalized by the mean intensity of the pre-bleaching frames. Another region outside the cells but close to the bleached region was selected as a background measurement. Using these regions, the bleach-corrected normalized mean intensity of the bleached region, *I*(*t*), was calculated as

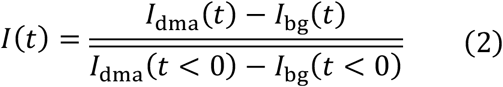

where *I*_dma_(*t*) is the spatial average of the bleach-corrected intensity of the bleached region, *I*_bg_(*t*) is the spatial average of the bleach-corrected intensity of the background region near the ER but where no cells were present. *t* < 0 indicates the pre-bleaching frames, and the overline indicates a time-average of the spatially-averaged intensities before bleaching. The recovery curve was fit to the model for a circular bleaching spot with a rectangular profile (30) (Fig. 2 B)

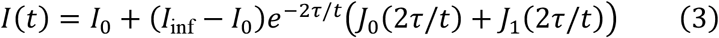

where *I*_0_ is the fluorescence intensity immediately after bleaching, *I*_*inf*_ is the fluorescence intensity long after recovery, *J*_0_ and *J*_1_ are the zeroth and first Bessel functions, respectively, and τ is the characteristic diffusion time, τ = *w*^2^/4*D*, where *w* is the radius of the bleaching spot and *D* is the diffusion coefficient.

#### PALM

CF488 was excited by 488 nm at 0.45 W/cm^2^ for 1 second to find the PM of cells and to draw a mask for patterned photoactivation (Fig 3 A, B and C). A patterned 405 nm laser was used to photoconvert mEos2 from the green state to the red state and a 561 nm laser was used to excite mEos2 in its red state (Fig 3 A and B). The illumination sequence consisted of one frame of 405 nm photoactivation with increasing power followed by nine frames of 561 nm excitation at a framerate of 20 Hz for up to 40,000 frames. Every 100 frames the 405 nm photoactivation frame was replaced by low power 488 nm excitation to potentially prolong the mEos2 trace lengths (31) and to potentially detect an increase in autofluorescence.

**Figure 3:**
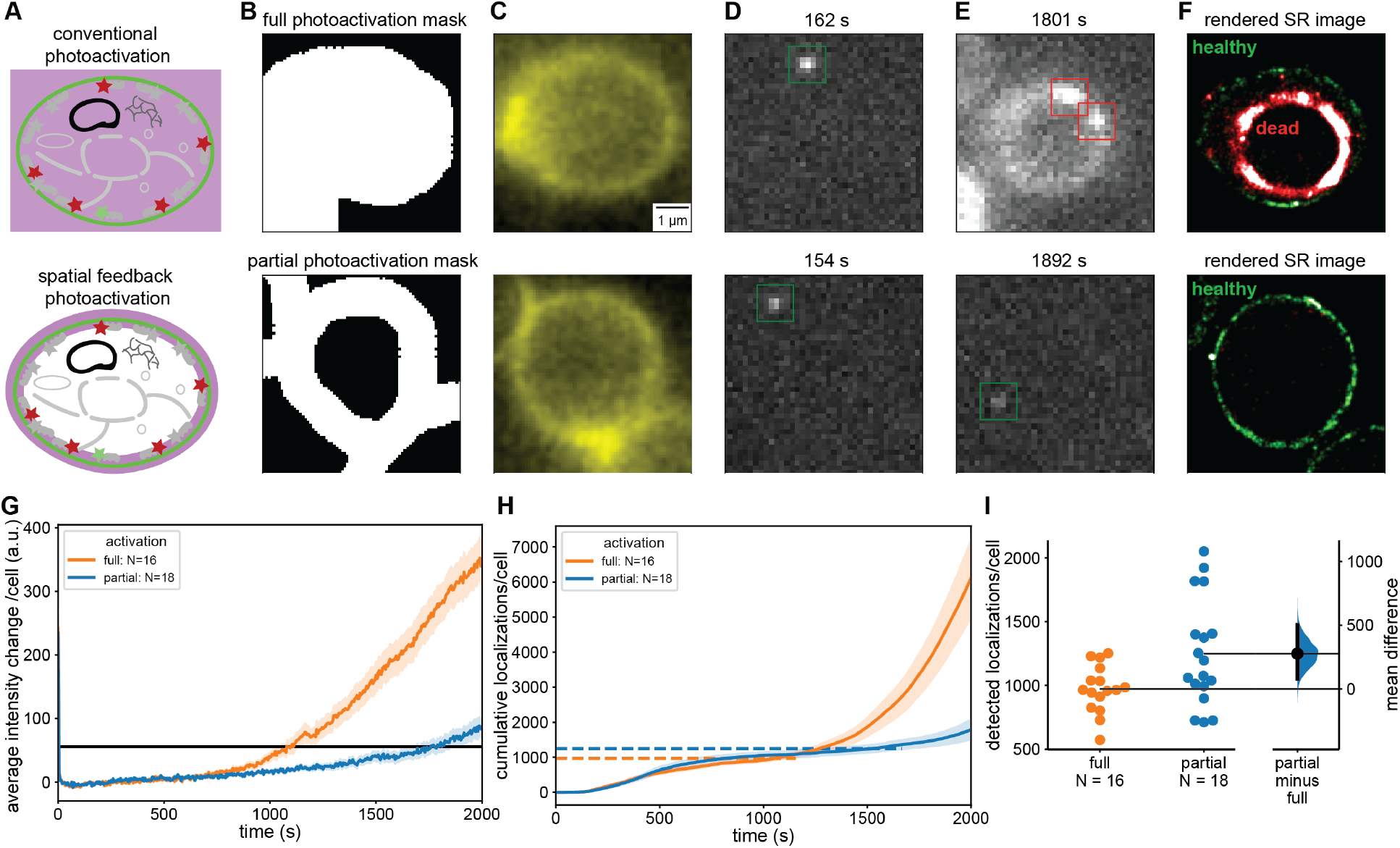
Comparison of conventional and spatial feedback PALM. (A) In conventional PALM the entire cell is exposed to 405 nm photoactivation light (upper) whereas in spatial feedback PALM only the regions that contain PAFPs are photoactivated (lower). (B) A mask is selected covering the entire cell (upper) and another mask which excludes the cytoplasm covering only the PM where mEos2 is located (lower). (C) Fluorescence image of CF488M conjugated to ConA excited at 488 nm used to create the mask in (B). (D, E) Individual frames with single molecule localizations of mEos2 under 561 nm excitation at early (D) and late (E) timepoints during PALM data acquisition. The fully illuminated cell shows signs of phototoxicity including autofluorescence much sooner (upper) than the partially illuminated cells (lower). (F) Rendered PALM image showing mEos2 localizations while cells are healthy (green) and after the transition to the unhealthy state (red). (G) Mean autofluorescence intensity change of fully and partially illuminated cells (orange and blue respectively) under 561 nm excitation and cutoff for transition to unhealthy state (black line). (H) Mean cumulative localizations of fully and partially illuminated cells (orange and blue, respectively) with horizontal dashed lines showing the mean number of localizations at the cutoff time. Localizations after the transition to the unhealthy state are unreliable and predominantly caused by autofluorescence. (I) Gardner-Altman plot of the total number of localizations from fully and partially illuminated cells up to transition to unhealthy state compared at the 95% confidence interval. (Mann-Whitney p < 0.05) (41) A significant 28% increase in reliable mEos2 localizations is detected in spatial feedback PALM compared to conventional PALM. Error bands: standard error of the mean across cells.

Single-molecule localization microscopy (SMLM) analysis was performed using Insight3 software (Zhuang lab, Harvard) (Fig. 3 D, E and F). Single-molecule recognition was confirmed by visual perception of fluorescent blinking and single-molecule identification parameters for 2D Gaussian point-spread functions (PSFs) were set accordingly (Gaussian height ≥ 47 photons, width 280-750 nm, ROI: 7×7 pixels). All single-molecule localizations were rendered as uniform Gaussian peaks. All super-resolution images were represented across at least 36,000 frames (Fig. 3 F and S7).

To account for differences in the initial intensity of each cell, the average intensity during the first 2001 frames was subtracted from the mean intensity trace of each cell (Fig 3 G).

The mean cumulative counts were calculated by using the ROIs around each cell for measuring the mean intensity but excluding the cytoplasmic background localizations that are more dominant in fully photoactivated cells. We then averaged the cumulative counts of localizations per cell at each frame across all fully and partially photoactivated cells (Fig. 3 H and I).

Since SMLM cannot be reliably performed when the background intensity of cells increases due to phototoxic effects, we determined a single cutoff intensity for all cells, at which the false positive rate of single-molecule localizations became excessive and at which cells transitioned to an unhealthy state. We determined the cutoff intensity change (56 counts) from a single movie by averaging the intensity change of three partially photoactivated cells with the lowest intensity in the last frame. Single-molecule localizations were considered invalid after the time at which each cell’s mean intensity change exceeded the cutoff intensity of 56 counts (Fig. 3 G, H, I and S4).

Single-molecule traces were made from localizations before the cutoff time. Using custom software written in MATLAB (The MathWorks, Natick, MA), single molecule localizations were linked within a radius of 0.9 μm and only traces with a length of at least 3 frames (0.15 seconds) were used for further analysis (Fig 4).

**Figure 4:**
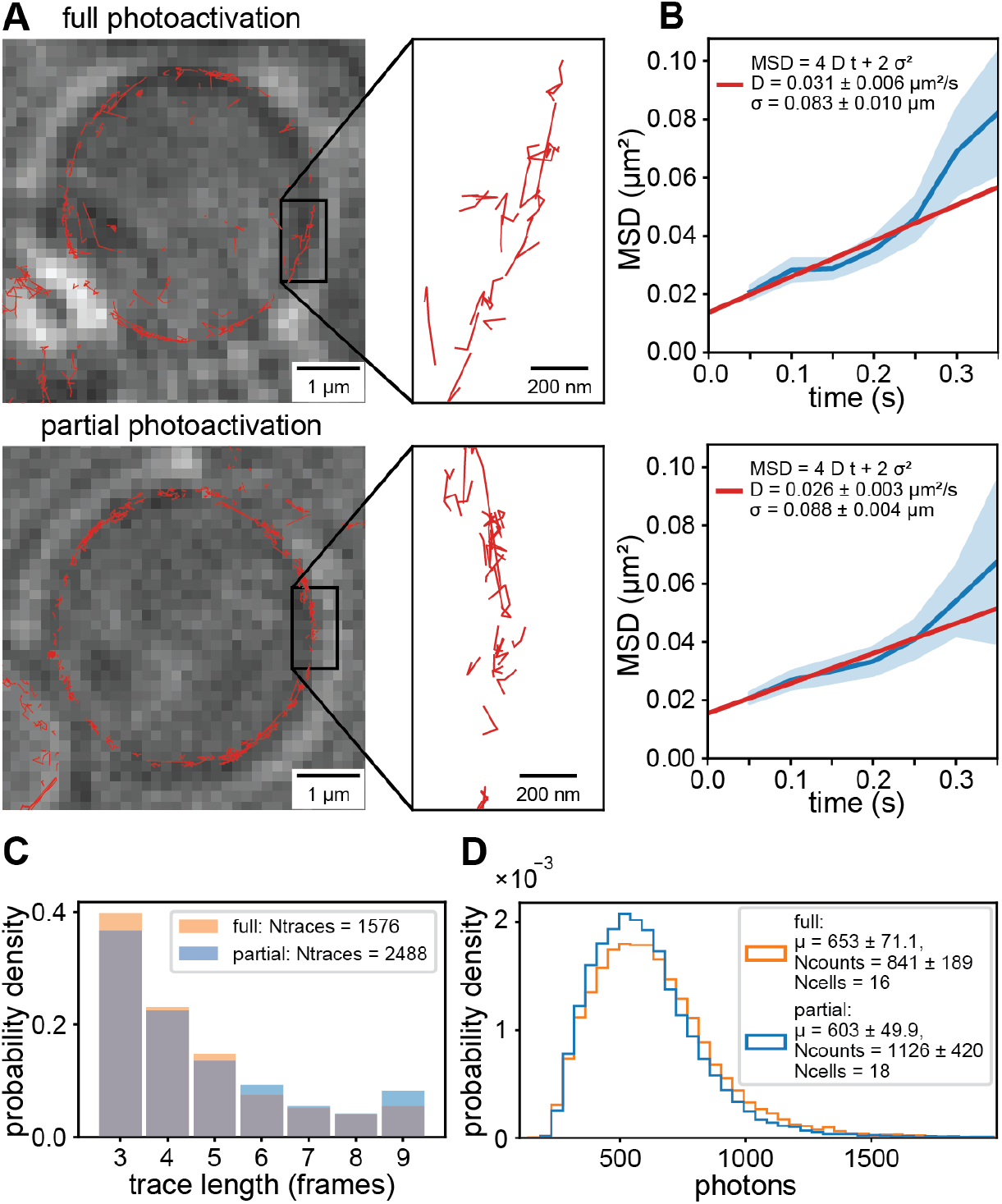
Comparison of single molecule tracking with conventional and spatial feedback PALM. (A) Single molecule traces (red) superimposed on LED images (average of 10 frames) of fully (top) and partially photoactivated cells (bottom). (B) Mean squared displacement vs. time (blue) and fit to model (red) of fully and partially illuminated cells (top and bottom, respectively). Error bands are the standard error of the mean across each lag time. (C) Probability density of trace lengths showing that the trace length statistics is the same for fully and partially photoactivated cells. All single molecule traces and localizations were analyzed up to the mean transition time to unhealthy state. (D) Probability density of photons per single-molecule localization showing that the quality of localizations is the same for fully and partially photoactivated cells.

To determine the diffusion coefficient, *D*, of mEos2 fused to the PH-domain of Plc(delta1) in fully and partially illuminated cells, we calculated the average mean squared displacement (MSD) from single-molecule traces by averaging over all squared displacements at each lag time, *Δt*. In the case of normal diffusion in two dimensions, the MSD is linear in time. We obtained *D* by fitting the MSD data to the model (32) (Fig. 4 B)

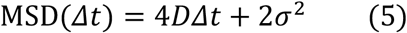

where *σ* is the uncertainty in position (32, 33).

## Results and discussion

### Computer-based calibration of a DMA allows for spatial light modulation with 52 nm precision

A general challenge for precisely modulating light in the sample plane of a microscope is to map pixels of a DMA to the camera pixels. While linear offsets in the sample plane can be manually corrected by adjustments, nonlinear distortions caused by the optical components in the excitation path remain to be a source of misalignment. Here, we develop a fast and simple computer-based calibration procedure, which accounts for both, translational misalignments as well as nonlinear distortions. First, a regular pattern of equally spaced spots is projected in the sample plane and the resulting fluorescence image is recorded with the camera. Next, the transformation between the coordinate system of the DMA and the sample plane is determined by fitting a 3^rd^-order polynomial to the centers of the spots. By applying the inverse transformation to any arbitrary pattern, all offsets and distortions are corrected, and the DMA and camera coordinate system are matched (see also Materials and Methods, Fig. 1 and Fig. S2).

We performed the calibration by projecting a 4×4 grid of spots, each with a diameter of 1.83 μm and a 9.53 μm center-to-center spacing, onto a layer of fluorescein in the sample plane (Fig. 1 A and Fig. S2). The projected pattern covered most of the field of view, which we restricted to 256×256 camera pixels, or 41×41 μm^2^. After determining the centroids of these spots by fitting each to a Gaussian profile, we calculated the inverse transformation that returns the original projected pattern. The 16 points were fit to a 3^rd^-order polynomial in two variables (20 parameters, see Eq. 1 and Materials and Methods). To demonstrate the advantage of our computer-based calibration we simulated a manual adjustment of the DMA by only allowing translation, scaling, and rotation about the beam axis in the transformation, which are the degrees of freedom that could be optimized by manual alignment. This transformation resulted in a significant discrepancy since it cannot correct nonlinear distortions of the optical system (Fig. 1 B). By calibrating with our computer-based approach and allowing skew degrees of freedom in the 1^st^-order and curved lines in higher orders of the transformation, a maximum deviation smaller than 52 nm was achieved in the 3^rd^-order transformation (Fig. 1 B and Fig. S2). This deviation is well below the optical diffraction limit and on the order of the single-molecule localization precision, which is tens of nanometers (9–12). Computer-based calibration is not only more precise than manual calibration but is also a fast and convenient way to account for small changes in the alignment of optical components. Performing a calibration before each experiment can be done in a total of less than twenty minutes including instrument startup and dye preparation. These results demonstrate that our computer-based DMA calibration is an easy and fast approach to precisely match the coordinate system of the DMA and the camera for advanced fluorescence microscopy applications with patterned illumination.

### Application of a calibrated DMA to Fluorescence Recovery after Photobleaching experiments

As a demonstration of this system’s accuracy for conventional microscopy applications, we performed a fluorescence recovery after photobleaching (FRAP) experiment with GFP fused to IRE1, which localizes to the endoplasmic reticulum (ER) of *S. cerevisiae*. In a FRAP experiment, a conventional fluorescence image is taken first to record the spatial distribution of the labeled molecules and to select a small region for photobleaching using a high-powered laser. A well calibrated DMA is therefore critical to precisely define the region of interest (ROI) for bleaching. This photobleaching is followed by a low-powered laser excitation of the entire field of view to measure the fluorescence recovery in the bleached region while the fluorescent membrane-bound molecules diffuse back. The diffusion coefficient of the mobile species present in the membrane can then be derived from the rate of fluorescence recovery and the fraction of mobile species can be determined from the steady-state intensity of the ROI after bleaching was applied.

When the diffusing protein, IRE1-GFP, was excited with a 488 nm laser at low power, the detected fluorescence signal showed the expected outline of the ER around the nucleus and in proximity to the plasma membrane of the yeast cells (Fig. 2 A, left). Due to the precise calibration of the DMA, the recorded fluorescence image allowed us to select a small region for bleaching on the DMA (Fig. 2 A, second from left). During bleaching, only the selected region of the DMA was turned on and the resulting bright fluorescence caused by the high excitation power confirmed the accuracy of the DMA calibration (Fig. 2 A, center, and Fig. S3). Due to the extension of the selected spot beyond the ER of the cell and due to optical diffraction, the shape of the fluorescent signal is not identical to the selected region. After bleaching the selected region, the entire field of view was excited again at low laser power to monitor the fluorescence recovery in the bleached region (Fig. 2 A, second from right and right). The normalized and bleach-corrected intensity of the fluorescence recovery curve in the region of the cell outlined in red (Fig. 2 A, second from right and right) allowed us to determine the diffusion coefficient of IRE1-GFP in the ER by fitting the model in Eq. 3 (Fig. 2 B). Based on the model, the diffusion coefficient was determined to be 0.00276 ± 0.00018 μm^2^/s with a mobile fraction of 89%. These results demonstrate that our calibration procedure enables accurate selection of regions from a microscopy image for FRAP experiments.

### Spatial feedback photoactivation enables longer imaging times in single-molecule localization microscopy with an improved number of localizations

One challenge in SMLM is the long imaging time caused by the requirement for sparse photoactivation and the need to acquire enough single molecule localizations to fulfill the Nyquist criterion for resolving a cellular structure. In addition, imaging for a long time can be necessary for studying dynamic processes in live cells such as observing the time evolution of intracellular signals or organelle trafficking. A major problem associated with long imaging times is the exposure of cells to the high energy photons from the 405 nm photoactivation laser, which causes phototoxicity in all cell types (34–36). Here we employ our calibrated DMA to develop a spatial feedback approach for photoactivation, which reduces phototoxic effects, results in more localizations compared to traditional photoactivation and extends the viable SMLM imaging time. In this approach a conventional fluorescence image is acquired to record the diffraction-limited spatial distribution of a protein of interest. Next, this image is used to create a binary mask on the DMA and to expose only those regions within cells to 405 nm photoactivating light where the proteins of interest are located. The entire field of view is then excited for nine frames with 561 nm light to image and localize single molecules. This cycle of spatially patterned photoactivation and excitation is repeated thousands of times until all labeled proteins are activated and imaged.

As a model system to demonstrate spatial feedback photoactivation for SMLM we used a *S. cerevisiae* strain in which the plasma membrane (PM) localizing PH domain of Plc(delta1) fused to the photoactivatable fluorescent protein (PAFP) mEos2 is expressed at low levels. While in principle the green pre-activated state of mEos2 could be used to create a conventional fluorescence image for the patterned photoactivation mask, we stained the cell wall in addition with fluorescently-labeled ConA, which is excited with low power of 488 nm light. This additional staining increases the fluorescence signal from the proximity of the PM and decreases bleaching of pre-activated mEos2 by requiring lower excitation powers. First, we excited the fluorescently-labeled ConA with a low power of 488 nm light and recorded a conventional fluorescence image of cells to create a photoactivation mask around the entire cell as a control for conventional PALM (Fig. 3 A and B, upper). Next, the mask was transformed with the inverse transformation between the camera and DMA coordinate system as previously described in the Material and Methods. A PALM data acquisition sequence consisting of one frame of 405 nm photoactivation followed by nine frames of 561 nm excitation for up to 2000 seconds. Every 100 frames the 405 nm photoactivation frame was replaced by low power 488 nm excitation to potentially prolong the trace lengths (31) and potentially detect an increase in autofluorescence. During the 561 nm excitation frames, initially single mEos2 molecules became visible at the PM (Fig. 3 D, upper) and were fitted to determine their precise location. However, after longer data acquisition, the autofluorescence of the cells started to increase, which indicated phototoxic effects (Fig. 3 E, upper). Localizations recorded after this transition to an unhealthy cell state were considered unreliable because of the cell’s altered physiological state and because of the high fluorescence background. After even longer imaging times, the cell’s morphology was altered (shrinking) and only bright autofluorescence was falsely detected and fitted as single molecules (Fig. 3 E, upper). The rendered super-resolution image in Fig. 3 F, upper, shows the expected outline of the PM for localizations before the transition to the unhealthy state (green). After the transition to an unhealthy state and the change in cell morphology only false localizations were detected inside the original outline of the PM (red squares in Fig. 3 E, upper).

In the same PALM movies, we randomly selected cells for spatial feedback photoactivation (Fig. 3 A, lower). A mask for photoactivation was created only around the PM of these cells where the PH-mEos2 protein is localized, leaving a region inside the cell without 405 nm light exposure (Fig 3 B, lower). The same PALM imaging sequence was used as before, and single molecules were fitted (Fig. 3 D and E, lower) and rendered in a super-resolution image (Fig. 3 F, lower). In strong contrast to the fully photoactivated cells, the partially activated cell did not show any signs of phototoxicity long imaging times and only some cells exhibited a slow increase autofluorescence towards the end of the PALM movie. Therefore, more reliable localizations were detected, and false positive localizations were almost negligible. Almost all localizations from the cell in Fig. 3 F, lower, show the PM localizations as expected and due to the longer imaging time contain more dense localizations compared to the fully activated cells.

To define the transition of each cell to an unhealthy state, we plotted their mean intensity change when excited with 561 nm light (Material and Methods and Fig. S4). As can be seen in Fig. 3 G, the mean intensity change of fully activated cells consistently increases after the transition from a healthy to an unhealthy state. The same trend can be seen in mean the cumulative number of localizations per cell (Fig. 3 H), which only contain false positives after the transition to an unhealthy state due to the bright fluorescence background. For each cell, we then defined the transition to a noticeable change in mean background fluorescence to be 56 counts (Fig. 3 G and S4 and Materials and Methods). While this threshold is somewhat arbitrary, it is the lowest that is visually noticeable in each frame and above the noise of the mean intensity vs. time traces of each cell (Fig. S4 A). Furthermore this threshold can be consistently applied to all fully and partially photoactivated cells. From the single-molecule data, fully illuminated cells showed an average increase in background of 121% after their respective cutoff times while partially illuminated cells only showed only a 47% increase (Fig. S5). The increase in autofluorescence background of fully illuminated cells therefore not only reports on the altered physiological state and morphological change of cells but also causes a high false positive rate of localizations and a high background level in those localizations (Fig. S5 and Fig. S7).

As indicated in Fig. 3 F and G, the mean time to reach an unhealthy state for partially photoactivated cells was 44% longer compared to fully activated cells and some partially activated cells even did not reach this state within the imaging time of 2000 seconds. We note that the mean intensity at the mean transition time for fully activated cells is lower compared to partially activated cells because some partially activated cells did not reach the intensity threshold within the imaging time. These results demonstrate that spatial feedback photoactivation allows for significantly longer imaging times. Importantly, this extended imaging time also significantly increases the number of localizations per cell and thus the quality of super-resolution images. By determining the number of localizations per cell until the transition to an unhealthy state, we found a significant 28% increase in the mean number of localizations from partially activated cells compared to fully activated cells (Fig. 3 I). While the distribution of the number of localizations is broad compared to fully activated cells, this result shows the recording SMLM data with spatial feedback photoactivation significantly increases detected localizations and improves the quality of super-resolution data.

### Feedback photoactivation results in the same localization precision and ability for single molecule tracking as conventional SMLM

Single molecule tracking gives valuable and complementing information about the dynamics of single molecules such as their diffusion coefficients, differences in mobility or transport modes (37). To verify that spatial feedback photoactivation does not alter the localization precision or the accuracy of single-molecule tracking compared to traditional SMLM, we performed single molecule tracking experiments with the same 2xPH-domain fused to mEos2 under partial and full photoactivation conditions. For both imaging modes, single molecule traces were created by linking localizations that appeared within 0.9 μm and within 1 frame (see Materials and Methods). Given our average photoactivation density of 0.003 localizations per frame and per μm^2^, the probability for accidentally linking two separate molecules is less than 0.3%.

Under both, full and partial photoactivation, we observed the expected traces of single molecules along the plasma membrane of yeast cells (Fig. 4 A). To further analyze the diffusion of molecules, we calculated the mean square displacement of traces from fully and partially photoactivated cells and fitted the linear model of Eq. 5 (see Materials and Methods). The resulting diffusion coefficients for full and partial photoactivation agreed within the error of the experiment (Fig 4B). Since the length of single molecule traces affects the accuracy of determining diffusion coefficients, we compared the histograms of trace length under full and partial photoactivation and did not detect significant differences (Fig. 4 C).

In SMLM, a larger number of photons emitted by a single emitter increase the localization precision as σ~1/√*N* (38). We therefore compared the photon statistics of reliable localizations for full and partial photoactivation. Again, no significant difference in the mean number of photons between fully and partially illuminated cells was detected (Fig. 4D). These results demonstrate that spatial feedback photoactivation has no effect on the ability to accurately localize single molecules and to study their diffusion in SMLM applications. The increase in imaging time due to reduction of phototoxicity as well as the increased number of reliable localizations therefore presents a significant advantage of spatial feedback photoactivation compared to the traditional photoactivation of the entire cell. Our simple and accurate calibration approach for patterned illumination applications as well as the development of spatial feedback photoactivation will therefore become of broad applicability that extends far beyond the proof of concept of this work.

## Conclusions

In this work we presented a fast, easy and accurate approach to calibrate DMAs for patterned illumination applications in fluorescence microscopy. Our computer-based transformation between the camera and DMA coordinate system replaces the need for small manual adjustments and results in an improved accuracy of 52 nm due to the additional nonlinear degrees of freedom that cannot be optimized otherwise. We demonstrated the usefulness of this approach in a conventional FRAP experiment with the ER localized protein IRE1 fused to GFP. Furthermore, we employed the accurate calibration of the DMA to develop a novel spatial feedback photoactivation approach for SMLM. In this approach only regions within the cell are exposed to 405 nm light in which photoswitchable fluorescent proteins are located. As a result, the possible imaging time until signs of phototoxicity were detected increased by 44% and 28% more single-molecule localizations were recorded. Spatial feedback photoactivation results in the same localization precision and ability for single molecule tracking as expected. Therefore, spatial feedback photoactivation presents a significant improvement of SMLM experiments compared to the traditional photoactivation of the entire field of view and will enable a myriad of applications in quantitative cell biology to study cellular processes and structures below the optical diffraction limit.

## Author Contributions

A.M. and E.M.P. designed approaches and experiments, interpreted data and wrote the manuscript. A.M. performed experiments, data analysis and programming. A.M. L.D. and C.T.E. designed, built and optimized the DMA excitation path.

## Acknowledgements

We thank Elizabeth M. Smith for making the IRE1-GFP yeast strain. Research reported in this publication was supported by the National Institute of General Medical Sciences of the National Institutes of Health under award number R21GM127965.

The authors declare no financial conflicts and interest.

